# Localizing metabolic synthesis in microbial cultures with kinetic mass spectrometry imaging (kMSI)

**DOI:** 10.1101/050658

**Authors:** Katherine B. Louie, Benjamin P. Bowen, Rebecca Lau, Trent R. Northen

**Affiliations:** Lawrence Berkeley National Laboratory, Berkeley, CA 94720, USA.

## Abstract

Mass spectrometry imaging (MSI) has emerged as a powerful technique enabling spatially defined imaging of metabolites within microbial biofilms. Here, we extend this approach to enable differentiation of newly synthesized versus pre-existing metabolites across a co-culture. This is accomplished by MS imaging two soil microbes, *Shewanella oneidensis* MR1 and *Pseudomonas stutzeri* RCH2, that were administered heavy water (D_2_O) during growth on agar plates. For two species-specific diglyceride (DG) lipids, isotopic analysis was performed on each spectra collected across the co-culture to determine the relative amount of newly synthesized versus pre-existing lipid. Here, highest levels of new synthesis of RCH2 lipid was localized to border regions adjacent to *S. oneidensis* MR1, while the MR1 lipid showed highest levels in regions further from RCH2. Interestingly, regions of high lipid abundance did not correspond to the regions with highest new lipid biosynthesis. Given the simplicity and generality of using D_2_O as a stable isotopic probe combined with the accessibility of kMSI to a range of MSI instrumentation, this approach has broad application for improving our understanding of how microbial interactions influence metabolite biosynthesis.

## INTRODUCTION

Soil microbes are continuously adapting metabolic processes in response to numerous stimuli, including changing microenvironment and nutrient availability. Presumably one of the greatest influences on microbial metabolism is the presence of other microbes in their environment^1^. Yet, how these complex communities are able to coordinate metabolism is poorly understood. This is largely because microbes are typically studied either in isolated cultures or complex natural environments (e.g. soils), making it difficult to identify the chemical responses arising from specific interactions between microbes.

Mass spectrometry imaging (MSI) has emerged as a powerful technique for deconstructing microbial interactions, making it possible to map metabolite location and relative abundance across an image. This field is developing rapidly and has been recently reviewed^2, 3^. Briefly, a broad range of mass spectrometry techniques including MALDI, DESI, nano-DESI, NIMS, and nano-SIMS have been used for creating molecular images of microbial isolates, co-culture and intact communities by rastering across a sample while simultaneously generating spatially defined mass spectra. In addition to detecting small metabolites and proteins, a more recent area of focus in mass spectrometry imaging has been the investigation of secondary metabolite production found in plant, fungal and microbial co-cultures^4-8^.

Recently, in moving beyond single static images and towards visualizing metabolic dynamics, time-course profiling during colony growth has been applied using MALDI-IMS^8^. Given the destructive nature of the MALDI mass spectrometry imaging technique and high vacuum requirement, this approach requires replicate cultures imaged at different timepoints to determine what metabolic changes occurred over time. Ambient mass spectrometry techniques such as DESI and Nano-DESI are relatively non-destructive and can be performed directly on microbial colonies at ambient conditions such that in principle, the same colonies can be imaged multiple times^9^. Unfortunately, the resolution of this technique is typically lower than MALDI and solvent contact with the sample may affect colony growth and interactions.

Previously, we developed a stable isotopic labeling approach with deuterium oxide to examine regional lipid flux across a biological sample with NIMS, kinetic mass spectrometry imaging (kMSI)^10^. Here, we investigate microbial interactions by applying kMSI to measure relative biosynthesis levels of species-specific lipids in a co-culture. In this workflow, interacting colonies are pulsed with deuterium oxide (Figure 1A), during which time deuterium is metabolically incorporated into newly synthesized compounds, such as lipids (Figure 1B). Following mass spectrometry imaging of the labeled colonies (Figure 1C), spectra from each image pixel is isotopically analyzed to determine the relative amount of newly synthesized versus pre-existing lipid that contribute to the composite spectra (Figure 1D). Resulting “kinetic” images based on isotopic ratios and labeled fractions enable detection of metabolite turnover largely independent of ion abundance and matrix effects.

**Figure 1.**
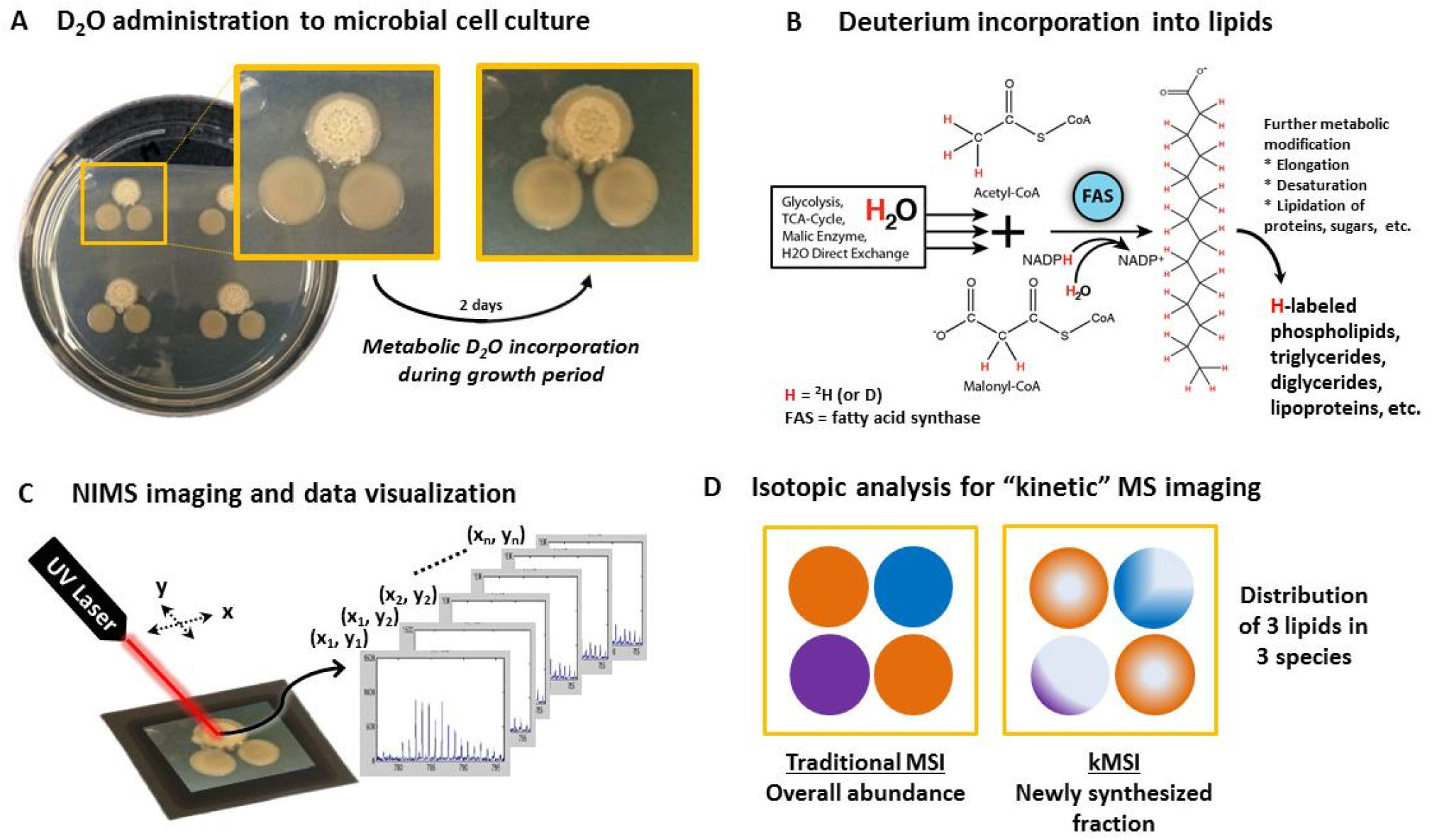
Workflow for “kinetic” imaging of microbial co-cultures. (A) Administration of deuterium during microbial co-culture to label newly synthesized compounds. (B) Metabolic incorporation of deuterium during lipid biosynthesis. (C) NIMS imaging of an isotopically labeled culture. (D) Comparison of ion intensity images in traditional mass spectrometry imaging versus “kinetic” MS images of metabolite synthesis based on isotopic analysis and deconvolution of labeled spectra for each image pixel.

This method was applied to examine the interactions of *Pseudomonas stutzeri* RCH2 and *Shewanella oneidensis* MR1, to examine the synthesis of two species-specific lipids. Using this kMSI approach, two species-specific lipids synthesized prior to and during the labeling experiment were found to be differentially located across the co-culture, with levels of new synthesis not reflecting that of lipid abundance.

## RESULTS AND DISCUSSION

Diverse metabolites play a wide range of activities in microbes ranging from biosynthesis and bioenergetics to complex chemical signaling^11^. It is well established that the dynamic production of many of these compounds occurs in response to the presence of and chemical signaling from other microorganisms^12^. Therefore it is desirable to have techniques that can both define the spatial localization and temporal dynamics of metabolism during colony interactions. Our previous efforts using D_2_O with LC/MS to investigate biofilm lipid dynamics and more recently kinetic mass spectrometry imaging of a tumor suggested that this approach may be suitable for imaging synthesis of lipid flux in a microbial co-culture as shown in Figure 1^10, 13^.

## Deuterium enrichment of microbial co-culture

Two bacteria, *P. stutzeri* RCH2 (RCH2) and *S. oneidensis* MR1 (MR1), were selected to test the kMSI approach for studying microbial colony interactions. Both are soil microbes easily distinguished by optical microscopy, having distinct color and morphology, as well as by the production of species-specific lipids detectable by mass spectrometry^14^. Liquid cultures were spotted on agar plates in a triangular pattern (2 spots of MR1 and 1 spot of RCH2) with each spot ^~^1 cm apart. After 2 days growth, the MR1 colonies appeared round, smooth and with a reddish hue, whereas the RCH2 colonies were whitish with a central rippled, rugose morphology, and beginning to grow towards MR1 (Figure 1A, left photo). At this point, a “pulse” of deuterium was added to the culture to isotopically label the metabolites being actively synthesized for the next 2 days of growth. This was accomplished by slicing out a small section of agar, filling the recess with 99.8% D_2_O that absorbed into the agar, then returning the culture to the incubator. Since the initial amount of water in the agar was ^~^10 mL, adding 800 µL D_2_O resulted in a final concentration of ^~^8% D_2_O. This percentage was used in the calculation of isotopic enrichment to model the isotopic composition of a fully labeled compound.

After the D_2_O pulse, the co-culture continued growing, with RCH2 moving even closer to MR1 and even starting to encroach upon the other microbe’s space (Figure 1A, right photo). During this interaction, since deuterium was simultaneously being incorporated into newly synthesized compounds, the metabolites actively involved could be identified by changes in isotopic pattern. To detect and visualize this isotopic incorporation, a mass spectrometry image was acquired of the culture using REX-NIMS^14^. Here, the colonies were dried and a methanol-soaked agarose gel was used to extract and transfer the metabolites from the colonies to the NIMS surface, analogous to liquid-liquid extraction commonly used for LC-MS. Importantly, this NIMS surface contained a previously spotted 400 µm × 400 µm grid of unlabeled MR1 extract to test the ability of kMSI isotopic analysis to differentiate ^2^H-labeled vs. the unlabeled lipids. Imaging was performed using 100 µm step-size resolution, acquiring ions over the mass range of *m/z* 50-1300 and viewed using OpenMSI^15^.

## Isotopic labeling detected in mass spectrometry image

Our analysis focused on two species-specific lipids that we characterized in previous work (*m/z* 523.5 and 575.5). These had been putatively identified as being structurally similar lipids but with different acyl chain lengths and degree of unsaturation, with the chemical formulas C_33_H_63_O_4_ and C_37_H_67_O_4_ based upon high resolution mass spectrometry and MS/MS showing fatty acyl chains with two fragments differing by 4H (corresponding to 2 double bonds) (Supplementary Figure 1, A-H)^14^. Initial work had putatively identified these lipids as quinones; however, further inspection of MS/MS spectra showed that the characteristic *m/z* 197 fragment ion of a ubiquinone headgroup was absent making a ubiquinone structure unlikely^16^ (Supplementary Figure 1, C-F). Instead, exact mass, chemical formula and fragmentation spectra for *m/z* 523.5 and 575.5 corresponded well with diglycerides, DG (30:0) and DG (34:2), respectively, ionizing as protonated species with water loss [M+H-H_2_O]^+^ (Supplementary Figure 1, C-H)^17^. However, we cannot be certain if these are DGs in the sample or DG fragment ions resulting from in-source degradation of phospholipids or triglycerides with loss of a headgroup^18^. DGs play many important roles in bacteria, ranging from modulating membrane fluidity in stress response, protecting against oxidative damage, cell-cell signaling, and also functional roles in lipoproteins, anchoring more hydrophilic proteins into a hydrophobic cell membrane^19-21^.

These two DG lipids were used to generate an image showing the spatial distribution of these lipids throughout the co-culture (Figure 2). Here, the spatial distribution of the MR1 lipid and RCH2 lipid in the mass spectrometry image correlate well with the colony locations in the corresponding optical image. Some distortion is apparent, however, which can be attributed to the dehydration step prior to metabolite extraction and transfer.

**Figure 2.**
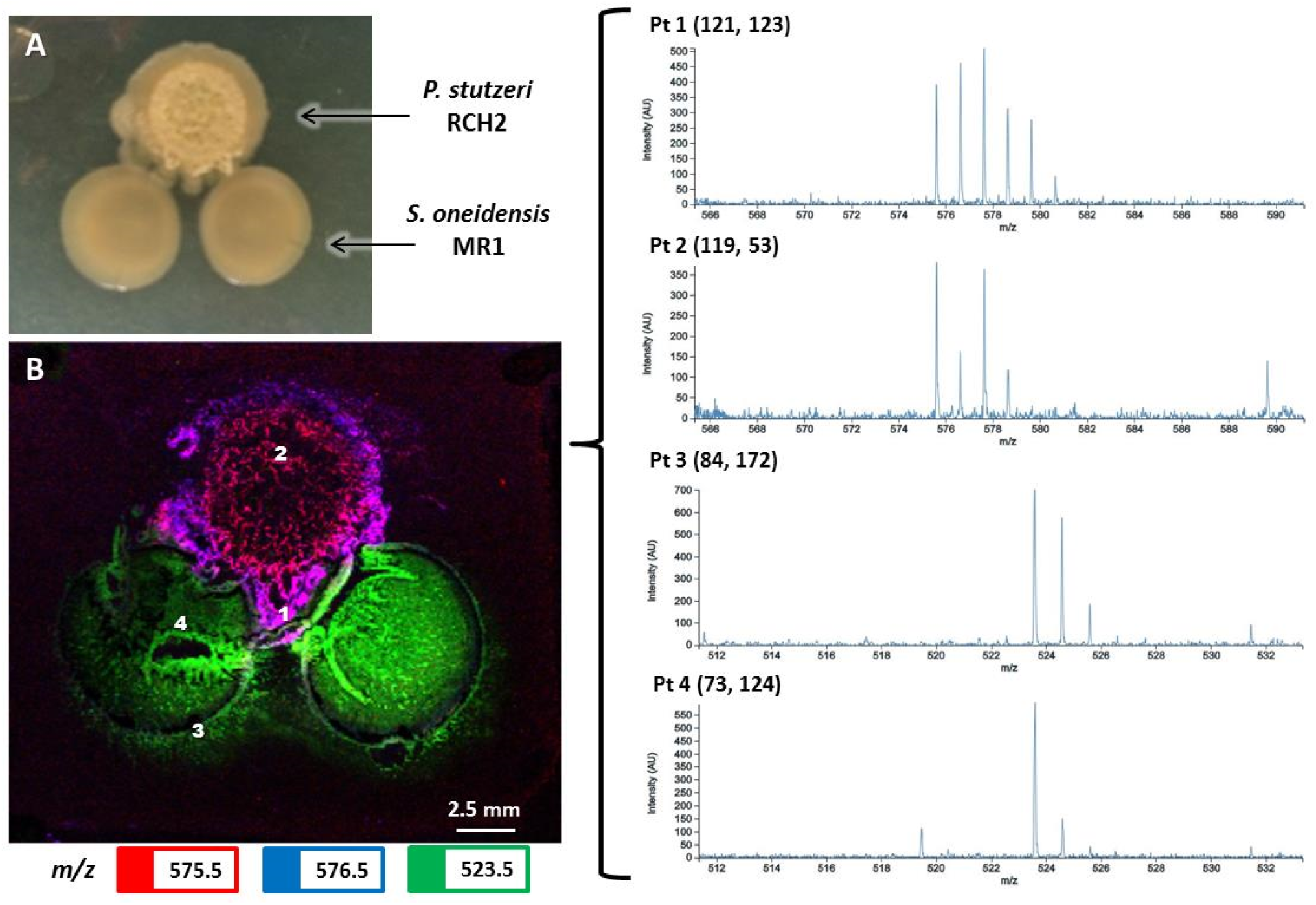
Visualization of differential isotopic labeling of species-specific lipids. (A) Corresponding optical image of *P. stutzeri* RCH2 and *S. oneidensis* MR1 co-culture imaged with kMSI. (B) Tri-color image of co-culture viewed in OpenMSI. Mass spectra corresponding to 4 different locations in the image show different levels of deuterium labeling for each species-specific lipid. *m/z* 523.5 for DG (30:0) from MR1 (Pt 3, 4) and *m/z* 575.5 for DG (34:2) from RCH2 (Pt 1, 2).

To confirm isotopic incorporation of deuterium, the isotopic patterns of the lipids were compared between pixels from different regions of the image. For the RCH2 lipid *m/z* 575.5 at point (119, 53), near the middle of the RCH2 colony, the M1 isotope is about 40% of the intensity of the M0 isotope. This ratio is comparable to that observed in the unlabeled spotted extract (Supplementary Figure 1B), indicating zero or only minimal labeling at this point. In contrast, at point (121, 123) near the colony edge, M1 is ^~^1.2x more intense than M0, indicating a large amount of isotopic incorporation of deuterium at this point. Differential labeling is also seen for the *m/z* 523.5 lipid, as exemplified in spectra from two pixels located either centrally or at the periphery of MR1 (Figure 2).

Here, it is important to note that changes in labeling pattern do not result from hydrogen-deuterium exchange since the solvent used in the extraction gel, methanol, is a protic solvent, not containing deuterium, and therefore any exchangeable sites on ^2^H-labeled compounds would become protonated during extraction. Thus, labeling patterns detected in the image were specific to only non-exchangeable locations, e.g. lipid backbones resulting from reduction reactions, and therefore could be attributed to active metabolic processes.

The overall regional distribution of labeling could be immediately visualized in the tri-color mass spectrometry image generated using the *m/z* 575.5, 576.6 and 523.5 ions as red, blue, and green, respectively. In regions where red is the dominant color, the intensity of the M0 isotope (red) of the RCH2 lipid is much higher than M1 (blue), indicating zero to minimal labeling; in contrast, in regions where a purple (red+blue) color dominates, the intensity of the M1 isotope increases relative to or exceeds that of M0, indicating larger amounts of labeling. While the red color is found centrally in the RCH2 colony, the color becomes increasingly purple towards the periphery, with large purple regions close to MR1. Since the purple color reflects higher amounts of labeling, this observation implies substantial new synthesis of the lipid in the RCH2 perimeter, especially near MR1. Overall, these gradations in color result from differential labeling patterns of the lipid throughout the image, which confirms that kMSI can visualize metabolic incorporation of deuterium, and provide insight into the different rates at which synthesis is occurring in across the colonies.

## Kinetic images of co-culture lipids

For the two diglycerides, DG (30:0) from MR1 and DG (34:2) from RCH2, the relative amount of newly synthesized versus pre-existing lipid was mapped throughout the image. This was accomplished by analyzing the isotopic pattern of each lipid in the spectra comprising each image pixel. To measure the maximally ^2^H-labeled isotopic pattern of the DG (30:0) lipid, MR1 was cultured in liquid media with 8% D_2_O and NIMS performed on MeOH extracts of the cell pellet (Supplementary Figure 2). The isotopic pattern for labeled DG (30:0) was used to determine a value for *N*, the maximum number of hydrogens in the molecule capable of being metabolically incorporated into a particular molecule originating from water (Supplementary Figure 3). The mass spectrum in each image pixel was then modeled as a composite of isotopic patterns from fully (8% D_2_O) ^2^H-labeled (at *N* locations) and fully unlabeled molecules, representing newly synthesized and pre-existing lipid populations, respectively, and deconvoluted to determine the relative contribution of each. This approximation of the isotopic patterns was used previously and enables the processing of large data files generated in mass spectrometry imaging^10^. This isotopic analysis is discussed in further detail in the Methods section – kMSI data processing.

By applying this analytical approach, the spatial distribution of the relative amounts of pre-existing versus newly synthesized DG (30:0) and DG (34:2) were mapped throughout the co-culture (Figure 3). Importantly, the acoustically printed grid of unlabeled MR1 extract confirmed the effectiveness of this approach in discriminating ^2^H-labeled versus unlabeled lipid populations since the grid appears only in the unlabeled image (Fig. 3D).

**Figure 3.**
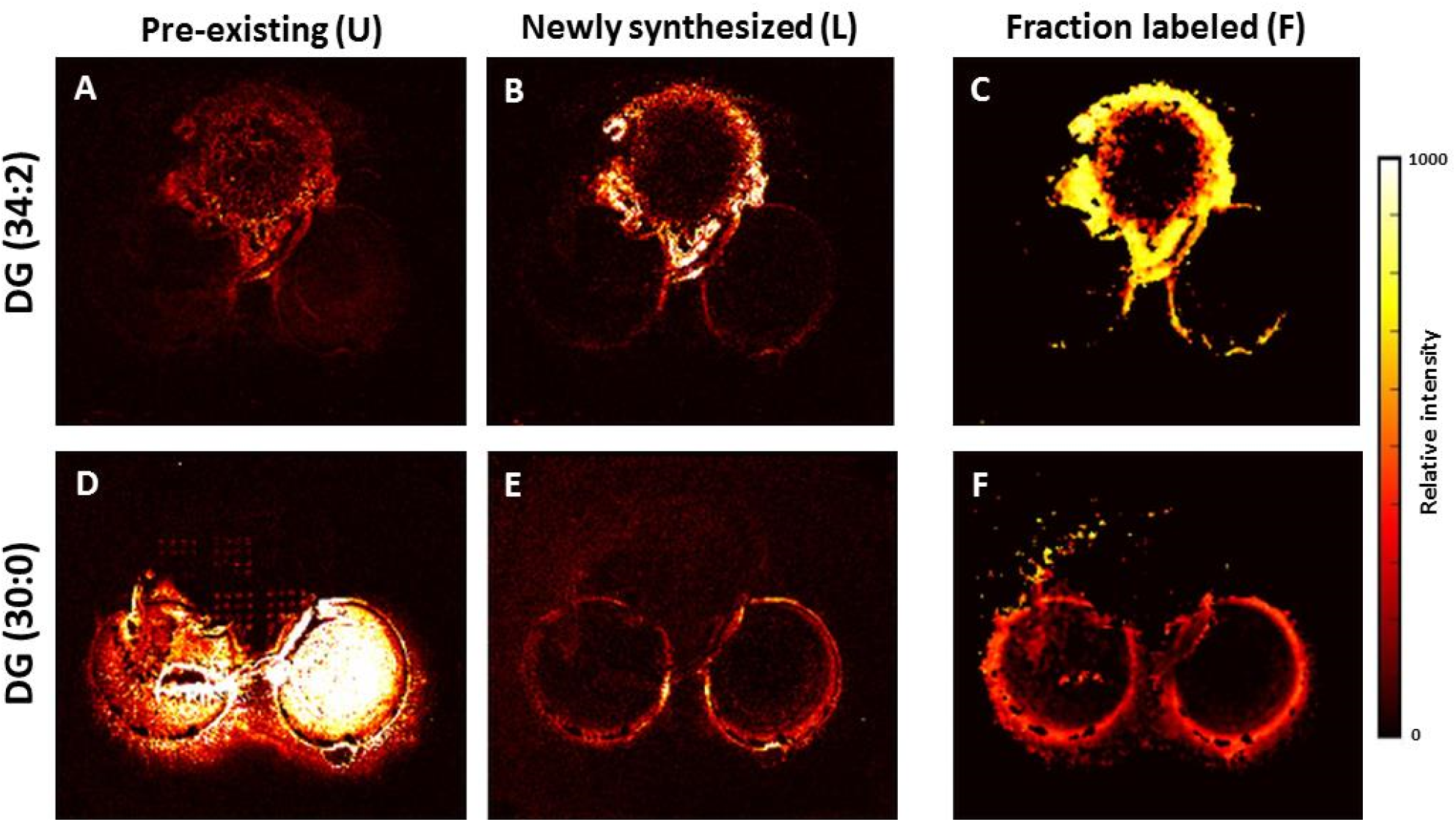
“Kinetic” mass spectrometry images for two lipids detected in co-culture showing the spatial distribution of DG (34:2) from RCH2 and DG (30:0) from MR1. Color scale indicates the relative level of each lipid: (A, D) Unlabeled, the pre-existing lipid population; (B, E) Maximally ^2^H-labeled, the newly synthesized lipid population and (C, F) Fraction ^2^H-labeled, the relative amount of lipid newly synthesized versus total. U=unlabeled, L=^2^H-labeled, F=fraction ^2^H-labeled=[L/(U+L)]

The kinetic images show that DG (30:0), although distributed throughout the two MR1 colonies and most abundant centrally (Fig. 3D), is being newly synthesized primarily at the perimeter of each colony (Fig. 3E), with the highest labeled fraction occurring at the edges furthest from RCH2 (Fig. 3F). In contrast, DG (34:2) from RCH2 shows a very different pattern, where the majority of both pre-existing and newly synthesized lipid occurs at the colony perimeter (Figure 3A, B), with larger relative amounts of new synthesis occurring in regions closer to the MR1 colonies and even beginning to encircle them (Fig. 3B, C).

The kinetic images show that high DG (34:2) intensity correlated with high levels of DG synthesis rates for RCH2, but for MR1, new DG (30:0) synthesis localized mainly in regions of low intensity. Examining the spatial patterning of DG (34:2) synthesis, we can speculate that it has a functional role in RCH2 movement towards MR1. This movement is apparent when looking at the optical images taken before (Fig. 2A, left) and after (Fig. 2A, right) deuterium administration. In addition, it has been shown in some Pseudomonas species that a lipoprotein lipid is used to tether pili used in motility to the outer membrane, so DG production may facilitate this function^22^. Alternatively, the increased synthesis of DG (34:2) may also be indicative of heterotrophy, with RCH2 utilizing a compound released from MR1, hence the movement towards the other microbe and increased regional synthesis. With MR1, again, we can only speculate as to why DG (30:0) synthesis decreased near RCH2. Possibly a secondary metabolite released by RCH2 is inhibiting growth of MR1; or, RCH2 is depleting a necessary nutrient and thereby limiting MR1 growth in the area. Although not conclusive, these observations from “kinetic” images provide a step towards understanding colony dynamics not available through static imaging.

The kMSI imaging approach for examination of microbial interactions has several advantages compared to traditional static mass spectrometry imaging. Just as stable isotope internal standards are valuable in quantification and normalization, images based on isotopologue ratios (e.g. Figure 3C, F) are less biased by matrix effects than those based on absolute ion intensity. For instance, across colonies, heterogeneity in surface pH, salt concentrations, etc. may favor ion detection in some regions while suppressing it in others, thereby confounding assertions based on relative levels of ion abundance^23^. In contrast, these factors do not bias kMSI images of labeled fractions since isotopologue ratio calculations do not depend on ion abundance. Though kMSI, like other isotopic tracing approaches, depends on uniform update of the probe, the high diffusivity and ubiquitous uptake of water by microbes suggest deuterium labeling should have minimal bias. The kMSI imaging facilitates the detection of regional metabolic flux within a single image taken at a single timepoint from a single sample. This effectively eliminates biological variation that may influence results when comparing replicate cultures imaged at different timepoints. Also, being largely independent of ion abundance, the technique is sensitive to small changes in biosynthetic levels and can be used to detect metabolite turnover even when metabolite utilization and synthesis occur at equivalent rates.

## CONCLUSION

Here we have applied kMSI to examine spatially defined lipid synthesis during growth of adjacent *Shewanella oneidensis* MR1 and *Pseudomonas stutzeri* RCH2 colonies. Our analysis effectively discriminated pre-existing vs. newly biosynthesized lipids based on the incorporation of deuterium added to the media during colony growth. While the highest levels of DG (34:2) production for RCH2 was adjacent to MR1 colonies, MR1 synthesis of DG (30:0) occurred primarily distal to RCH2. Unlike typical mass spectrometry imaging, relative abundance levels did not need to be compared to observe colony dynamics and matrix effects were minimized since images were based on ratios of labeled and unlabeled compounds. Overall, kMSI provides a valuable temporal dimension to MSI with potential to provide insight into dynamic biochemical processes to complement existing approaches.

## METHODS

### Chemicals

Chemicals used were LC-MS grade methanol (Sigma), Luria-Bertani (LB) Miller agar (J104, Amresco, Solon, OH), R2A media (M1687, HiMedia, Kennett Square, PA), MilliQ water, agarose (A9539, Sigma), bis(hepta decafluoro-1,1,2,2-tetrahydrodecyl) tetramethyl-disiloxane (Gelest, Morrisville, PA) initiator, and deuterium oxide (D_2_O, 99.8%, DLM-4-99.8, Cambridge Isotope Laboratories, Inc., Tewksbury, MA).

### Microbial cultures and deuterium administration

Microbial strains *Shewanella oneidensis* MR1 (ATCC Cat#700550) and *Pseudomonas stutzeri* RCH2 (a strain isolated from Hanford 100H, WA^24^) were cultured in liquid R2A media for two days at 30 °C from frozen glycerol stocks. For mass spectrometry imaging, 5 µL droplets of each microbe were spotted onto LB agar in a round 100mm dish. After two days of culture at 30 °C, ^~^1.3mL volume of agar was cut out with a sterile spatula to create a reservoir into which 800 µL deuterium was added. The D_2_O soaked into the agar, and the agar culture was further incubated at 30 °C for an additional two days, and subsequently prepared for mass spectrometry imaging. To prepare cultures with full deuterium labeling, additional cultures of MR1 and RCH2 were prepared in liquid media amended with either 8% D_2_O, then cell pellets extracted with MeOH after two days culture for NIMS analysis.

### Replica-extraction transfer MSI (RexMSI)

Preparation for imaging microbial cultures using replica extraction transfer for mass spectrometry imaging (RexMSI) on a NIMS surface has been thoroughly described^14^ as well as NIMS wafer fabrication^25^. Here, a section of agar containing spotted *P. stutzeri* RCH2 and *S. oneidensis* MR1 was excised and dried at 35 °C for ^~^2 hours on a glass slide in a food dehydrator (FD-61 Snackmaster, NESCO/American Harvest, Two Rivers, WI). Replica extraction transfer stamping of biomolecules from the microbial culture onto a NIMS surface coated with bis(heptadecafluoro-1,1,2,2-tetrahydrodecyl) tetramethyl-disiloxane initiator was then performed using a 4% agarose gel soaked in methanol.

### Acoustic printing

Prior to RexMSI of the microbial culture, the NIMS chip was spotted with an MR1 extract. Here, an acoustic liquid transfer system (ATS-100 Acoustic Liquid Dispenser, EDC Biosystems, Fremont, CA) was used to print 10 nL droplets of microbial extract reconstituted in 60% acetonitrile in a 400 µm center-to-center grid pattern. Details of this process have been described previously^26^.

### NIMS imaging

Mass spectrometry image acquisition was performed using an AbSciex 5800 MALDI TOF/TOF (AbSciex, Foster City, CA) in positive reflector MS mode equipped with an Nd-YAG laser (200 Hz). Mass spectra were acquired over a mass range of 50-1300 Da while accumulating 15 shots/spot. The 4800 Imaging Tool software (Novartis and Applied Biosystems) was used to raster the laser across the NIMS surface and record spectra in an x-y step-size of 100 µm × 100 µm. MS and MS/MS fragmentation spectra from extracts of RCH2 and MR1 cell pellets were obtained using a MALDI LTQ Orbitrap (Thermo Scientific, Waltham, MA) in positive mode, acquiring full range MS spectra as well as MS2 via collision-induced dissociation (CID) and higher energy collision dissociation (HCD) on selected ions.

### kMSI data processing

Imaging files (.img) were uploaded into OpenMSI (https://openmsi.nersc.gov) for data visualization^15^. Relative levels of newly synthesized metabolites (labeled with deuterium, or ^2^H-labeled) versus pre-existing metabolites (unlabeled) were calculated based on a model of isotopic enrichment described in previous work with imaging deuterium-labeled lipids^10^. Here, model isotopic patterns of enrichment were generated for the molecules detected with *m/z* 523.5 and 575.5; these molecules were characterized as structurally similar diglyceride lipids, DG (30:0) and DG (34:2), respectively, with differing acyl chain lengths and number of double bonds based on exact mass and MS/MS fragmentation spectra. Based on this information, a chemical formula was assigned to *m/z* values 523.5 and 575.6 of C_33_H_63_O_4_ and C_37_H_67_O_4_, respectively, as well as 577.6, C_37_H_69_O_4_, assuming 577.6 to also be a DG but with 1 less double bond in the fatty acid chain giving rise to a 2 Da mass difference (DG 34:3).

Since the final measured mass spectrum in each image pixel is a composite of isotopic patterns from the detected ^2^H-labeled and unlabeled molecules, each spectrum was deconvoluted to determine the relative contribution of each. For an unlabeled lipid, the isotopic pattern was modeled based on chemical formula with natural distribution of heavy isotopes; for a labeled lipid, the isotopic pattern was modeled based on chemical formula with increased levels of the ^2^H isotope on a specific number of sites on a molecule. Thus, the maximum level of increase was determined by 2 criteria: (1) *D*, the % deuterium oxide in the agar and (2) *N*, the maximum number of ^2^H atoms capable of being metabolically incorporated into a particular molecule originating from water^27^. Here, the value of *D* was calculated to be ^~^8 atom% (800 µL of D_2_O was added to absorb into agar already containing ^~^10 mL water). The value used for *N* was 29 for *m/z* 523.5; this was experimentally determined by the isotopic pattern detected on NIMS resulting from spotted cell extracts of MR1 cultured with 8% D_2_O (Supplementary Figure 2, 3). Based upon this experimental result and previously reported values in literature for fatty acid chains on lipids, values of *N* used for 575.6 and 577.6 were 32 and 34, respectively, assuming *N_(523.5)_* +/-3d +/- 1.5s (where d is the number of incremental 2-carbon units on the fatty acid chain, and s the number of double bonds) to account for elongation and unsaturation^27^. These model parameters were then used to calculate relative levels of newly synthesized and pre-existing DG (30:0) and DG (34:2) lipids detected at *m/z* 523.5 and 575.6 for each image pixel.

## ACKNOWLEDGEMENT

This material by ENIGMA- Ecosystems and Networks Integrated with Genes and Molecular Assemblies (http://enigma.lbl.gov), a Scientific Focus Area Program at Lawrence Berkeley National Laboratory is based upon work supported by the U.S. Department of Energy, Office of Science, Office of Biological & Environmental Research under contract number DE-AC02-05CH11231.

## SUPPORTING INFORMATION

Further information can be found in a separate file - Supplementary Figures 1-3. This material is available free of charge via the Internet at http://pubs.acs.org.

